# A single complex V2 as an alternative model for the organization of mouse visual cortex

**DOI:** 10.1101/2024.10.10.617533

**Authors:** Declan P. Rowley, Mary Beth Cassity, Jimin Park, Ryan Gillis, Julien Corbo, Casey Lennon-Jones, Severine Durand, R. Carter Peene, Avalon Amaya, Hannah Belski, Ivana Bussi, Tye Johnson, Jaimie Kenney, Henry Loeffler, Robyn Naidoo, Ben Ouellette, Alex Stearns, Jackie Swapp, Madeline Tom, Julie Weber, Josh Wilkes, Allison Williford, Shiella Caldejon, Peter A. Groblewski, Jérôme A. Lecoq, Madineh Sedigh-Sarvestani

## Abstract

The mouse has become a major model for studies of the visual system, creating a need for principles of areal delineation that can be compared across species. In many well-studied mammals, V1 is bordered by a single second visual area, V2, whereas the cortex immediately surrounding mouse V1 is commonly parcellated into roughly eight higher-order visual areas (HVAs), with boundaries defined in part by reversals in the progression of the visual field. Work in other species has shown that such reversals can occur within a single visual area when retinotopic mapping is complex, raising the possibility that some conventionally defined mouse HVAs may instead be subdivisions of a single complex V2. We use simulations to show that complex retinotopy within a single visual area can generate multiple apparent areal borders beyond mouse V1. We also show that distinct biases in visual field representation among areas is a contributing factor to many of the reported functional differences among HVAs. Finally, our unified-V2 hypothesis makes several testable predictions, one of which is that functional differences between conventional HVAs need not exceed the variation within a single area. To test this prediction, we collected electrophysiological and two-photon recordings across V1’s retinotopic map, providing a large-scale dataset that we are making publicly available with this study. Using this dataset, we examined response properties associated with visual cortical hierarchy and found that they clearly distinguished V1 from higher visual cortex, while variation among individual HVAs did not significantly exceed variation across sampled locations within V1. Together, these findings support the unified-V2 hypothesis as an alternative model of mouse higher visual cortex and emphasize the importance of converging topographic, functional, architectonic, and connectional evidence in defining cortical areas.

---

What constitutes a cortical area is a central question in the study of cortical organization and function. This is not merely a question of nomenclature: areal parcellations guide hypothesis formation, experimental targeting, and interpretation of regional differences in computation. The criteria used to define areal boundaries therefore directly shape how cortical organization is studied and understood. Here we propose a re-evaluation of the current definition of these criteria and their application in the mouse visual cortex.

In the visual cortex, where retinotopy provides a measurable organizing principle, topography has served as a central criterion for area delineation. Garrett et al. (2014) formalized four retinotopic criteria for identifying mouse visual areas: uniform visual field sign within an area, absence of redundant representation within an area, redundant representation between neighboring same-sign areas, and reproducibility of area location across experiments. More generally, however, topography is only one source of evidence for areal boundaries, alongside functional specialization, architectonics, and connectivity. Evidence for a distinct area is strongest when multiple independent properties exhibit reproducible and spatially coincident transitions. Because functional, architectonic, and connectional properties can themselves covary with retinotopy, apparent convergence across modalities should be treated as independent evidence only when it cannot be explained by a shared topographic organization.

In the mouse, one of the main model organisms in neuroscience, the parcellation of higher order visual cortex has evolved in recent decades (Glickfeld & Olsen, 2017) with different studies parcellating the cortex beyond V1 into between 2 (Rose, 1929) and 16 areas (Zhuang et al., 2017). Projection and functional mapping studies provided increasingly detailed evidence for multiple retinotopic subdivisions beyond V1, establishing the parcellation that now dominates the field (Garrett et al., 2014; Wang & Burkhalter, 2007; Zhuang et al., 2017). Reversals in progression of the visual field, together with connectivity and cytoarchitecture, have played an important role in identifying areal boundaries (Figure 1A). Under a simple retinotopic mapping scheme, a reversal marks the beginning of a new map and therefore provides strong evidence for a new visual area.

**Figure 1.**
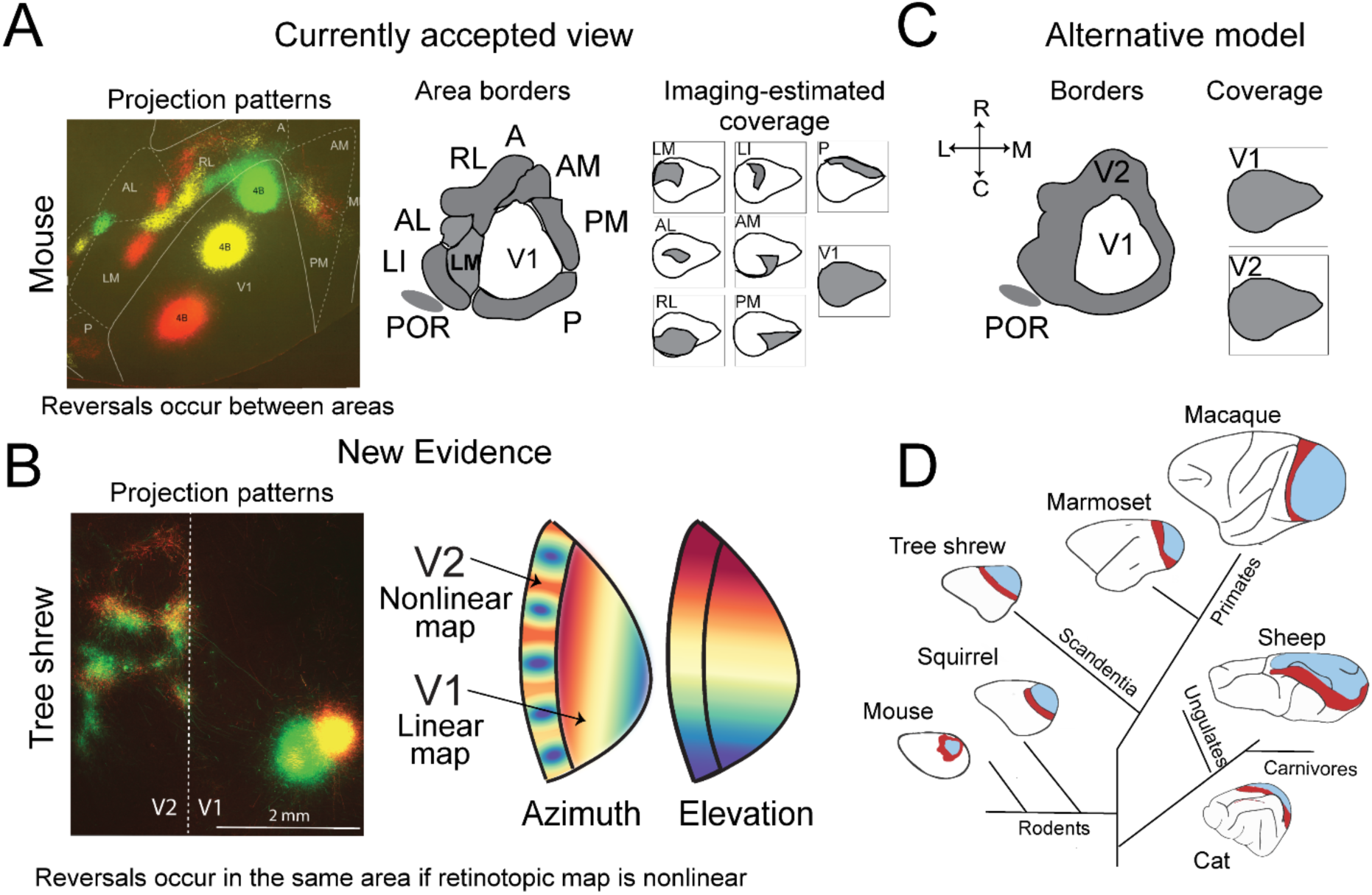
Unified-V2: A single complex V2 as an alternative organization of mouse higher visual cortex. **A)** The currently accepted organization of the mouse visual system includes multiple higher order visual areas beyond V1, delineated using functional retinotopy together with anatomical connectivity. Individual HVAs exhibit overlapping but systematically biased representations of visual space. Projection pattern figure is modified from (Wang & Burkhalter, 2007) and visual field coverage maps are modified from (Zhuang et al., 2017). These imaging-based maps may underestimate absolute visual field extent, but they reveal systematic differences in the distribution and weighting of visual space across HVAs. **B)** New evidence of visual field reversals within single areas in the tree shrew (Sedigh-Sarvestani et al., 2021) demonstrates that reversals do not necessarily indicate areal boundaries. **C)** We consider an alternative organization wherein many of the conventionally defined HVAs are subdivisions of a single area V2 in the mouse. **D)** Such an organization would bring the criteria used to delineate HVAs into closer alignment with those applied in several other species. Modified from (Rosa & Krubitzer, 1999).

Comparative evidence, however, shows that this inference does not hold universally. In many well-studied mammals, including primates, cats, and tree shrews, V1 is bordered by a single second visual area, V2, with additional higher-order areas lying beyond it (Rosa & Krubitzer, 1999) (Figure 1D). Moreover, visual field reversals can occur *within* individual visual areas such as V2 when the retinotopic transformation is complex. Such complex maps have been reported in tree shrews (Sedigh-Sarvestani et al., 2021), marmosets (Yu et al., 2020), ferrets (Manger et al., 2002), and squirrels (Kaas et al., 1989), and can arise from wiring-minimization constraints imposed by the geometry of cortical tissue (Wolf et al., 1994).

These observations suggest that retinotopic criteria must be interpreted in the context of cortical geometry and related complexity of the retinotopic map. In a simple retinotopic map, a repeated representation encountered after progression through visual space provides strong evidence for a second map belonging to a distinct area. In a complex map, however, the retinotopic transformation can fold or reverse direction within a single continuous area, producing both field-sign reversals and local redundancy in visual field representation (Figure 1B). Visual field reversals therefore cannot distinguish a boundary between two maps from an internal fold within one complex map. Comparative work has consequently raised the possibility that repeated extrastriate modules may sometimes represent subdivisions of a single V2 rather than separate higher visual areas (Kaas et al., 1989; Malach, 1989; Rosa & Krubitzer, 1999).

These considerations prompted us to ask whether some of the higher order areas of the mouse that are delineated primarily by visual field reversals, may in fact be subdivisions of a single area V2 (Figure 1C). Such a unified-V2 model could help explain the systematic and spatially organized biases in visual field representation reported across higher visual cortex (Figure 1A, Zhuang et al., 2017): Nearby HVAs in cortical space exhibit biased representation for nearby regions of the visual field, an organization consistent with a single area V2 exhibiting a complex retinotopic map. A unified-V2 would also bring the criteria used to classify mouse higher visual cortex into closer alignment with those applied to visual areas in other mammals (Figure 1D).

One challenge to this interpretation is functional specialization. Conventionally defined mouse HVAs differ in spectral sensitivity, spatial-and temporal-frequency preference, receptive-field size, response latency, binocular disparity, and other functional properties (Glickfeld et al., 2013; Marshel et al., 2011; Rhim et al., 2017; Sit & Goard, 2020). These differences are important evidence for functional organization, but they need not necessarily imply distinct areas. Because mouse HVAs also differ systematically in the portions of visual space they emphasize, some functional differences could be associated with biased sampling of an underlying retinotopic organization within a single area (Sedigh-Sarvestani & Fitzpatrick, 2022).

Here we use computational modeling and reconsideration of published data to explore the unified-V2 hypothesis as an alternative organization of mouse visual cortex. Under the unified-V2 model, many conventionally defined HVAs may instead constitute spatially organized, functionally distinct subdivisions of a single complex V2. We ask whether a single retinotopic map constrained by cortical geometry can generate the reversals and projection patterns used to delineate HVAs, and to what extent functional differences among these regions may reflect their biased sampling of visual space rather than discrete areal identity. In addition, we release a new large-scale dataset comprising electrophysiological and two-photon measurements across multiple positions within V1. Using V1 as a known single-area benchmark, we ask whether functional differences used to characterize hierarchy among HVAs exceed the variation that can occur across different locations within one visual area. Together, these analyses introduce and explore the unified-V2 model, while providing an open dataset for evaluating various models of mouse visual cortex organization.

## RESULTS

### Complex retinotopy within a single visual area can generate the appearance of multiple higher order areas in mouse visual cortex

The cortex immediately adjacent to mouse V1 was originally designated cytoarchitectonically as areas 18a (lateral) and 18b (medial) and early electrophysiological mapping identified a single area V2 within lateral 18a, bordered by V3, together with two smaller maps in medial 18b, as reviewed in (Glickfeld & Olsen, 2017). Subsequently, precisely localized multi-color projection studies (Wang & Burkhalter, 2007) revealed multiple topographically organized projection zones beyond V1. Subsequent functional retinotopic mapping identified multiple field-sign reversals and redundant representations, providing the basis for the HVA parcellation widely used today (Garrett et al., 2014; Zhuang et al., 2017). However, as discussed above, multiple visual field reversals can be a hallmark of complex retinotopic mapping within a single area (Sedigh-Sarvestani et al., 2021; Yu et al., 2020). Inspired by foundational theoretical work showing that cortical geometry can generate retinotopic discontinuities within a single map (Wolf et al., 1994), we set out to explore the reversals beyond mouse V1 with an elastic net model of retinotopy, well-established for simulating cortical map organization in other species (Goodhill & Willshaw, 1990; Sedigh-Sarvestani et al., 2021; Wolf et al., 1994; Yu et al., 2020).

Here, we model the cortical region surrounding V1 as a single continuous area. For simplicity, we call this simulated region V2. Its cortical extent is constrained by the V1 border and a population-derived estimate of the surrounding extrastriate cortical footprint from Waters et al. (2019), rather than by the boundaries of individual HVAs. This simulated V2 consists of point “neurons,” each with a receptive field (RF) in the visual field. The objective of the algorithm is to arrange the RFs so they tile the visual field with the wiring-length constraint of preserving local map smoothness, agreement with prescribed V1 border RFs and local isotropy. The algorithm iteratively optimizes the RFs to reduce the total cost associated with visual field coverage and smoothness over a fixed optimization schedule.

Figure 2 shows the outcome of an example instance of the model simulating the mouse visual system in V2, or the cortical space beyond V1 using 2,560 neurons. A single elastic net model, constrained by the geometry of V2 and trained to cover the mouse visual field, can generate retinotopy with reversals similar in number and scale to those reported in the mouse. This builds on earlier theoretical work showing that complex and fractured retinotopic maps, including visual field reversals, can emerge within a single area as a consequence of constraints imposed by cortical geometry (Wolf et al., 1994). Here, the reversals are likewise not imposed on the model; they emerge from the competing constraints of visual field coverage, local map smoothness, and the geometry of mouse visual cortex. Using matched parameters, the model produced a relatively simple V1 map, when constrained by the geometry of V1, and a more complex map with internal reversals, when constrained by the ring-like geometry of V2 surrounding V1 (Figure 2A-B). The precise locations of these reversals vary with model parameters and initial conditions and are not intended as an exact fit to the empirical HVA map. Rather, the simulation provides proof of principle that an explicitly defined single visual area can develop a complex retinotopic map containing multiple field-sign reversals when constrained by the ring-like geometry of mouse cortex beyond V1. Thus, the model demonstrates the feasibility of multiple reversals arising as structure within a single complex map, without imposing internal areal boundaries.

**Figure 2.**
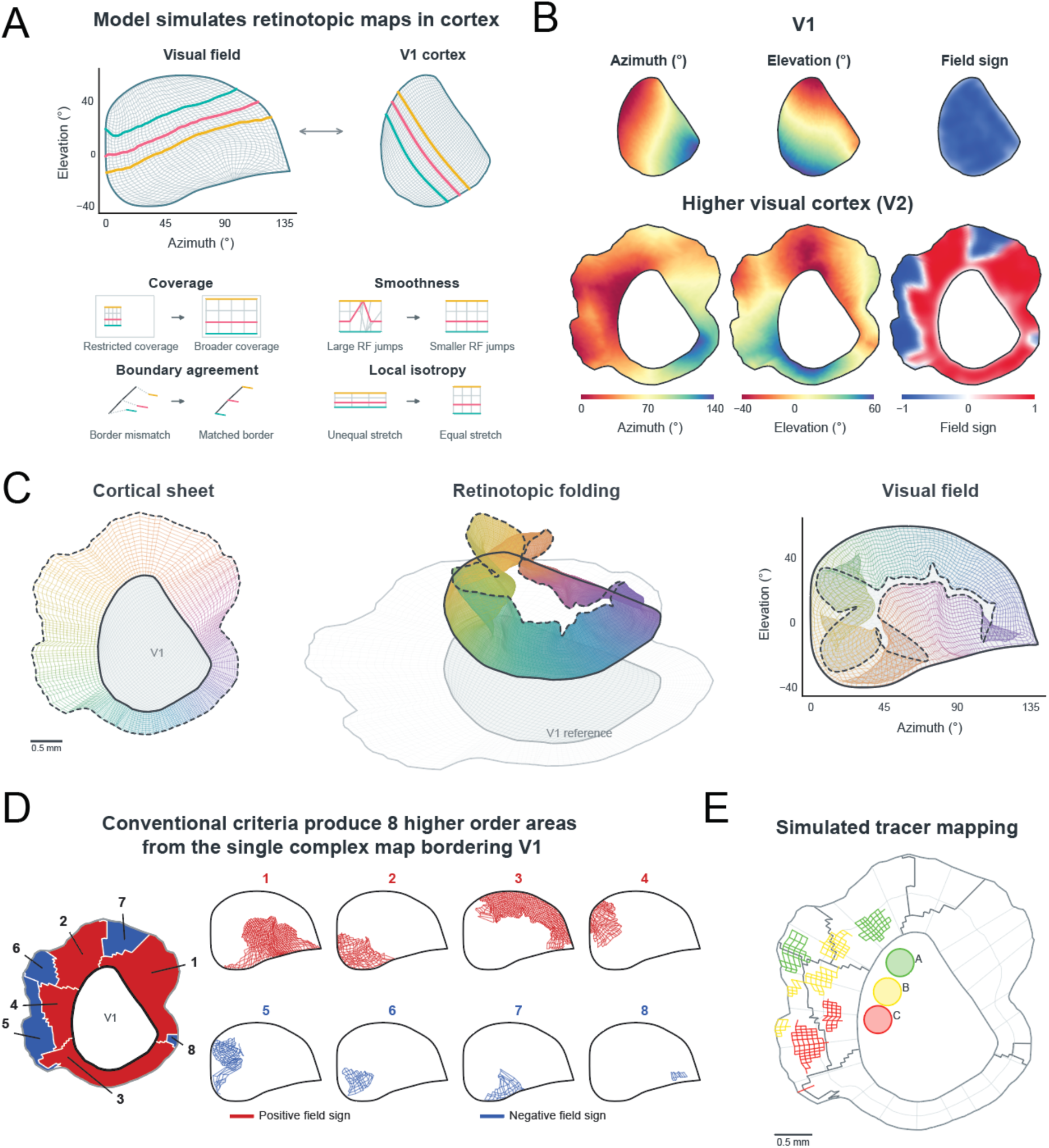
Retinotopic models show emergence of a single complex map of visual space beyond mouse V1. We extended the elastic net model from previous work (Sedigh-Sarvestani et al., 2021) to constrain a model of V2 to the unique structure of the mouse. **A)** Fitting the same model to V1’s cortical footprint results in a simple map with a 1-to-1 mapping between cortex and visual field. Colored transects identify corresponding locations. Schematics illustrate the objectives of the model: coverage of the visual field, local smoothness, continuity with the fixed border RFs and local isotropy. **B)** Azimuth and elevation are smooth across the V1 map and the field sign (Sereno et al., 1994) is uniformly negative, indicating a continuous mirrored map of the contralateral visual hemifield. The same model, when applied to a cortical footprint derived from Waters et al. (2019) for the region of cortex surrounding V1 results in a complex retinotopic map with non-monotonic progression, retinotopic reversals, evident in the alternating field sign. **C)** The higher visual cortex is visualized in cortical coordinates (left), over the corresponding retinotopically matched cortical coordinates in V1 (middle) and in visual field coordinates (right). Height in the middle plot is an illustrative coordinate based on the cortical distance from the border, illustrating the idea that the surface is more constrained near the V1 border due to the influence of the boundary agreement constraint. The folds reflect reversals in the visual field progression within the single modeled area. **D)** Applying an adaptation of the area-delineation rules of Garrett et al. (2014) to this single map yields eight candidate ‘areas’, four positive-sign (red) and four negative-sign (blue), each with a distinct bias in visual field coverage. **E)** Simulated projections from 3 spatially segregated patches within V1, along with the resulting projection pattern of connections beyond V1, illustrating how patchy projection patterns in retinotopically matched regions of visual space of V1 and a single higher order area can emerge from a complex retinotopic transform. The visual field for the model was implemented as a smooth envelope constrained by electrophysiological estimates of V1 coverage from Wagor et al. (1980; approximately 0–140° azimuth and −40–60° elevation) and the rounded coverage reported by Garrett et al. (2014).

The difference between the V1 and V2 maps can be understood by comparing the shapes of their cortical surfaces with that of the visual field (Figure 2A–B). A retinotopic map assigns each location on the cortical sheet to a corresponding location in the visual field. In V1, both surfaces are simple, filled-in shapes, allowing a smooth, one-to-one mapping between them, producing a simple retinotopic map without reversals (Figure 2B, top). In contrast, the cortical region surrounding V1 is ring-shaped, with a hole occupied by V1 itself (Figure 2B, bottom). In topological terms, a surface with a hole cannot be mapped continuously and one-to-one onto a surface without one. When the model attempts to represent the same visual field on this ring-shaped cortical sheet, while preserving local smoothness and continuity with V1, the resulting retinotopic maps often fold back on themselves, producing multiple reversals (Figure 2B, bottom).

These folds are illustrated in Figure 2C, where the modeled V2 cortical sheet (left) is plotted over corresponding retinotopic locations in V1 (middle). An artificial vertical coordinate represents distance from the V1 border, allowing the progression of the map to be followed as the cortical sheet extends outward. The resulting surface folds over itself, reflecting reversals in visual field progression and producing overlapping representations of visual space (Figure 2C, right). These are not physical folds in the cortex, but folds in the retinotopic mapping. Thus, a single continuous modeled area can exhibit both field-sign reversals and locally redundant representations of visual space (Figure 2D), features conventionally used to delineate separate mouse HVAs.

Partitioning the model’s single retinotopic map for the cortical sheet around V1, according to visual field sign produces, eight apparent areas, each with a different bias in its representation of visual space (Figure 2D). These biases are not independent: neighboring cortical regions tend to represent neighboring portions of the visual field. This organization resembles the orderly differences in visual field representation reported across mouse HVAs (Wang & Burkhalter, 2007; Garrett et al., 2014; Zhuang et al., 2017). As our model makes clear, such a systematic relationship between visual field bias and cortical position across regions is consistent with their being subdivisions of a larger continuous retinotopic map.

We next used the model to simulate the consequences of tracer injections into V1, assuming projections from V1 target retinotopically corresponding locations in higher visual cortex (Figure 2E). Simulated injections at three locations in V1 produced multiple spatially segregated projection patches beyond V1, despite the modeled cortex being a single continuous area. Although the model is not intended to reproduce the precise projection patterns reported by Wang & Burkhalter (2007); it demonstrates that patchy projections patterns can emerge from a single complex retinotopic map. Together, these simulations show that field-sign reversals, locally redundant visual field representations, systematic differences in visual field coverage, and segregated projection zones are all compatible with a single complex visual area. The model does not establish that mouse higher visual cortex is a unified V2, but demonstrates that these features are not, on their own, sufficient evidence for multiple distinct areas.

### Functional organization across higher visual cortex is strongly structured by retinotopy

Functional specializations are one of the main metrics used to justify areal definitions in the higher order visual system of primates and cats. For instance, area MT in primates exhibits enhanced sensitivity to global motion direction relative to area V2, even in identical regions of the visual field. Such hierarchical and parallel processing of overlapping parts of the visual field is thought to represent an efficient strategy to build selectivity to complex input features (Felleman & Van Essen, 1991). However, established visual areas may also contain specialized subdivisions. Macaque V2 stripes process overlapping portions of the visual field and demonstrate reproducible differences in their feature tuning (Hubel & Livingstone, 1987; Livingstone & Hubel, 1984). Functional differences alone are therefore not sufficient to define an area, and differences that simply track visual field position are even weaker evidence, since they are expected within any single retinotopic map. The relevant question is therefore whether the functional differences among mouse HVAs indicate distinct specializations warranting separate areal identities, or reflect retinotopic or modular variation within a larger area.

Under the unified-V2 hypothesis, conventionally defined HVAs are considered functional modules of a single area, and their response properties need not be uniform. Functional differences could take two forms. They could reflect specialized modules within the area, as with the stripes of macaque V2. Or they could follow visual field position, in which case they should vary smoothly across the cortical sheet, including across borders defined by retinotopic reversals, rather than changing abruptly at conventional areal boundaries.

Functional differences that follow visual field position, and therefore do not argue for separate areas, could arise from at least three non-mutually exclusive sources. First, the differences may already be present across the retinotopic map of V1 and simply conveyed to retinotopically matched locations in V2. Second, they may be delivered by subcortical pathways from the superior colliculus and the lateral posterior nucleus (LP). These regions project to higher visual cortex with strong retinotopic biases and functional response properties that differ across those biases (Tohmi et al., 2014; Bennett et al., 2019; Juavinett et al., 2020; Calzoni et al., 2026). Third, they may be generated within V2 itself, through target-specific selection of V1 inputs (Glickfeld et al., 2013) or through local circuitry. The first two sources predict that functional differences among mouse HVAs should largely follow visual field position and extend smoothly across conventional areal boundaries. We test this prediction first against existing observations and then by reanalyzing a published dataset.

In the mouse higher order cortex, many reported functional properties vary continuously across multiple conventionally defined areas, rather than forming robust and sharp differences between areas as observed in primates (Figure 3). For instance, binocular disparity tuning (La Chioma et al., 2019), color tuning (Aihara et al., 2017), and coherent motion processing (Sit & Goard, 2020) have all been shown to vary across the elevation axis of retinotopy (Figure 3A). Cortical inputs to upper-and lower-visual field domains of the superior colliculus are strongly organized by visual field position across extrastriate cortex (Calzoni et al., 2026). These trends are consistent with the ethological relevance of visual field locations for a low-lying animal like the mouse (Qiu et al., 2021). Furthermore, the areas categorized into the dorsal and ventral streams of the mouse cortex also exhibit robust biases for the lower and upper visual field (Figure 3B). Functional property differences in higher order visual areas have also been reported to vary along the azimuth axis of retinotopy. Encoding of various behavioral task-related features was shown to vary across higher order visual areas of the mouse (Minderer et al., 2019), but as the authors point out, this variation is agnostic to higher order areal boundaries and instead follows the underlying topography smoothly (Figure 3C). In fact, the only functional discontinuities in the mouse posterior cortex were found at the major anatomical borders between V1, parietal, and retrosplenial areas. Finally, learning-related plasticity in the mouse visual system has been reported across broad medial and anterior higher-visual-cortex domains rather than individual named areas (Zhong et al., 2025). Together, these findings suggest that functional organization in higher visual cortex largely follows retinotopic or mesoscale cortical axes that traverse conventional HVA boundaries, rather than necessarily being organized into discrete area-specific domains.

**Figure 3.**
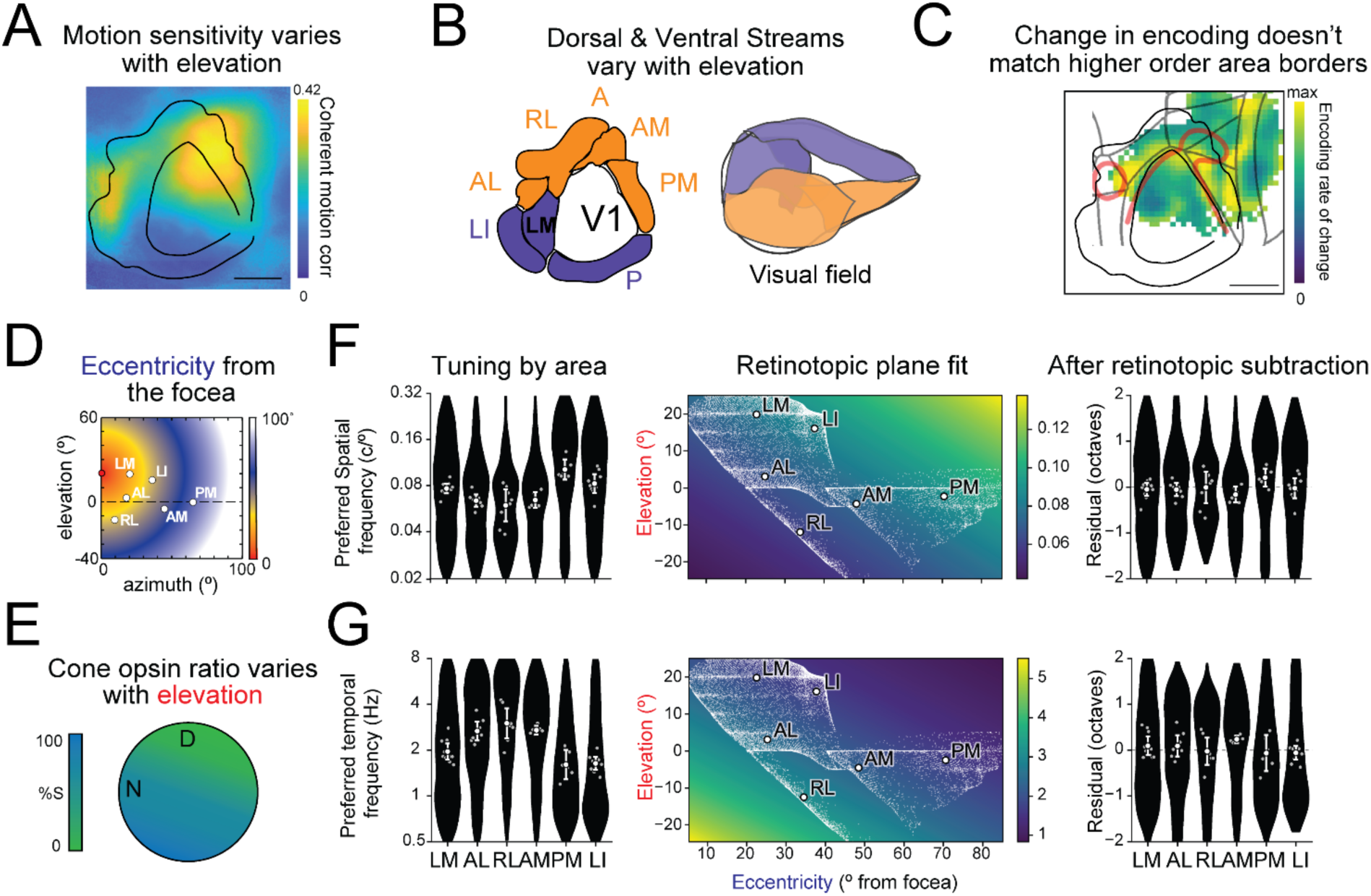
Retinotopic position contributes to functional variation across mouse higher visual cortex. **A)** Map of motion sensitivity adapted from (Sit & Goard, 2020) showing higher sensitivity in the lower field representation of V1 and higher order visual areas. **B)** Dorsal and Ventral streams in the mouse containing multiple higher order visual areas have been suggested due to differences in function, however these maps closely align onto elevation axis of retinotopy. Figure made using data from Zhuang et al. (2017) **C)** Behavioral encoding as a function of cortical space shows large change along the V1 border, but not between higher order visual areas (Minderer et al., 2019). Scale bar 1mm. **D)** Schematic of eccentricity from focea with area estimates for the data used in F&G. **E)** Schematic of retinal cone opsin gradient following (Demb & Singer, 2015). **F–G)** Reanalysis of Han et al. (2022) examining spatial-and temporal-frequency tuning before and after accounting for estimated retinotopic position. F) Preferred spatial frequency (SF). Left, cell-level tuning distributions across six higher visual areas. Middle, fitted SF as a function of elevation and eccentricity from the focea. Small white points indicate estimated retinotopic positions of individual cells included in the fit; large white points indicate the mean retinotopic position of the cells displayed for each area. Color indicates fitted SF. Right, residuals after subtracting the fitted retinotopic component; the dashed line indicates zero. Gray points show individual-mouse means, and large white points and bars show the equally weighted mean across mice and its 95% confidence interval. G) Same as F for preferred temporal frequency (TF). Plane fits included censored values at the analysis bounds; distributions and residual summaries include cells within those bounds (see Methods).

If functional differences among HVAs partly reflect which portions of visual space they disproportionately sample, it is important to distinguish visual field extent from visual field bias. Electrophysiological recordings (Wagor et al., 1980; Wang & Burkhalter, 2007; Wang et al., 2011) indicate that some HVAs span broader regions of visual space than suggested by widefield imaging, whereas functional imaging reveals substantial overlap together with strong differences in the relative weighting of visual field locations (Garrett et al., 2014; Zhuang et al., 2017). Importantly, these biases are spatially organized, such that neighboring HVAs preferentially represent neighboring portions of visual space (Figure 1A). Thus, our argument does not require incomplete visual field coverage: even if individual HVAs ultimately prove to span the full visual field, their systematically different weighting of central, peripheral, upper, and lower visual space could produce substantial differences in population-averaged response properties. Uncertainty in absolute extent therefore does not remove the orderly retinotopic biases most relevant to our functional argument.

Such observations raise the possibility that some functional differences among mouse HVAs are organized according to their systematically different weighting of the retinotopic map rather than discrete areal identity (Groen et al., 2022; Kanwisher, 2001; Sedigh-Sarvestani & Fitzpatrick, 2022; Yu & Rosa, 2014). Testing this hypothesis requires careful measurements of visual field location and functional property differences in V1 and higher order visual areas. Such data sets are sparse but growing. To test this retinotopic explanation with existing data, we reanalyzed a dataset from Han et al. (2022) and found that accounting for eccentricity and elevation reduced some of the reported differences in spatial-and temporal-frequency tuning among HVAs (Figure 3). The relationship of tuning with eccentricity is unexpectedly inverted relative to the classic trend reported in primate visual cortex, but is nonetheless still smooth.

In primate vision research ‘eccentricity’ is used to mean distance in visual degrees from the center of gaze, which is a useful metric for animals with a fovea and robust overrepresentation of the central few degrees of vision. Many functional properties scale with eccentricity within visual areas, making it a very important parameter to control for in foveal animals such as primates (Yu et al., 2015). However, in the afoveate mouse, ‘center of gaze’ is difficult to control for and typically elevation above eye level and azimuth relative to a normal through the eye are used. While this is a reasonable approach for an animal without a fovea, a central region of the upper binocular visual field of the mouse, named the ‘focea’, has been reported to have enhanced spatial resolution: V1 population receptive fields are smallest there, single-neuron receptive fields in LM and AL are smallest there, and behavioral acuity is highest there (van Beest et al., 2021). Notably, this region is not associated with increased cortical magnification in V1; the enhanced resolution reflects reduced receptive-field scatter and overrepresentation of binocular space rather than a foveal-like expansion of the map (van Beest et al., 2021). The focea nonetheless provides a natural reference point for a mouse analog of eccentricity. Taking the focea as 20 degrees above resting eye level on the vertical meridian (0 degrees azimuth), we defined ‘eccentricity’ (Figure 3D) as the Euclidean distance from this point in degrees of visual field, and assessed functional properties as a function of this distance.

There is an additional effect across retinotopy in the mouse that must be considered. The mouse retina has a gradient of cone opsin expression (Figure 3E) such that S-opsin, sensitive to UV light, is overrepresented on the ventral surface (the upper visual field) and gives way to M-opsin, sensitive to green light, on the dorsal surface (the lower visual field). When a typical LCD monitor is used, relatively more M-cones, compared to S-cones, are partially activated (Rhim et al., 2021). This means that retinal regions in the upper and lower visual fields are in a rod-dominated scotopic and mixed cone-and rod-mediated mesopic regime respectively. This retinal gradient is known to be preserved in the visual cortex (Rhim et al., 2017) and is thought to play an important role in predator evasion (Qiu et al., 2021). Thus, control for the opsin gradient is crucial for accurate measurement, reporting, and comparative analysis of functional properties in mice. Specifically, the use of typical LCD monitors results in primacy of scotopic vision in the ventral retina, producing higher spatial and lower temporal frequency tuning in the upper visual field.

In Han et al. (2022), an LCD screen was used. The observed trend, albeit small, toward higher SF tuning in areas with predominantly upper-field representations (Figure 3F) suggests that the cone-opsin gradient may contribute. To estimate the contribution of retinotopic position, we registered individual cells to the Zhuang et al. (2017) azimuth and elevation maps and fit SF and TF preference as a function of elevation and eccentricity from the focea (Figure 3F–G; Methods). Subtracting the fitted retinotopic component reduced some of the differences in mean SF and TF tuning among higher order areas. This suggests that visual field position contributes to measured functional differences among HVAs. We emphasize that this analysis is agnostic to the source of the retinotopic component it removes: the fitted plane describes how tuning varies with retinotopy across higher visual cortex, regardless of whether that variation is inherited from V1, from subcortical regions, or generated within V2. It also leaves open residual differences not captured by retinotopy, or those with more complex nonmonotonic relationships with retinotopy.

We note that changes in spatial and temporal frequency, varying smoothly with eccentricity, have been observed in visual areas of various species including primates (Sedigh-Sarvestani & Fitzpatrick, 2022; Yu et al., 2010). However, in the case of spatial frequency, the relationship we observe across mouse higher visual cortex appears to be inverted from that observed in primate V1 (Yu et al., 2010). In our reanalysis of Han et al. (2022), preference for higher spatial frequencies increases with eccentricity from the focea rather than decreasing (Figure 3F). We are not aware of a prior report of this relationship in mouse higher visual cortex; Han et al. (2022) and Marshel et al. (2011) reported area-level SF differences without analyzing SF as a function of retinotopic position, and Marshel et al. (2011) restricted their comparisons to eccentricity-matched neurons near the central visual field. Such an inverse relationship does not alter the evidence that SF preference is associated with retinotopy, although it changes how we interpret the source of these differences. The few available measurements within V1 show smaller receptive fields and higher behavioral acuity for the focea, the central region of the upper visual field, and a cutoff SF that increases with elevation (X. Zhang et al., 2015; van Beest et al., 2021). Taken at face value, this would suggest that the SF gradient across V2 modules is not simply inherited from V1. However, the evidence is sparse, the V1 and HVA measurements come from different stimuli and methods, and the cone-opsin gradient confounds both under conventional display conditions. Resolving this requires functional properties measured as a function of receptive-field position across V1 and across higher visual cortex under matched conditions; the dataset released here provides the V1 half of that comparison.

### Large-scale V1 recordings provide a within-area benchmark for functional heterogeneity

To more directly assess how much functional variation can occur within a single cortical area, we collected a new large-scale dataset through the Allen Institute OpenScope program. The dataset includes electrophysiological recordings from four targeted V1 locations per mouse as well as two-photon imaging recordings from 12 targeted V1 locations per mouse (Figure 4A-B).

**Figure 4.**
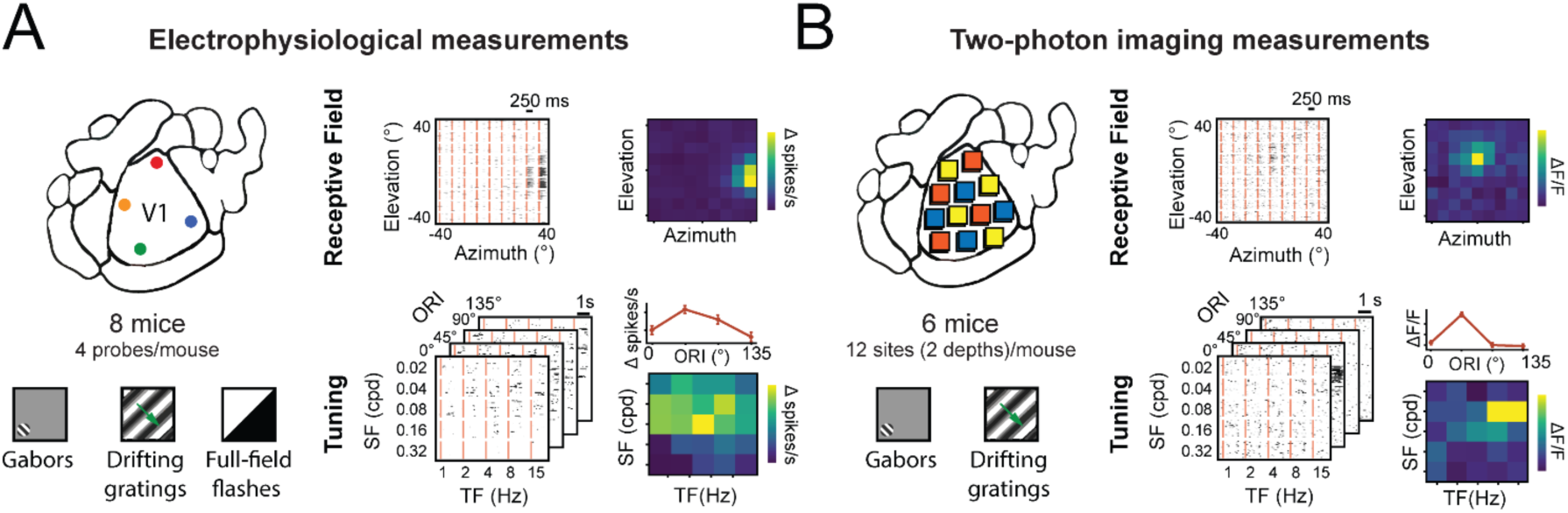
A new V1 dataset provides a within-area benchmark for functional variation across mouse visual cortex. This enables testing aspects of the unified V2 hypothesis. **A)** Electrophysiological recordings sampled four locations within V1 using Neuropixels probes, in each of eight mice. Visual stimuli included localized Gabors for spatial receptive-field mapping, full-field drifting gratings for characterization of orientation (ORI), spatial-frequency (SF), and temporal-frequency (TF) tuning, and full-field flashes. Example stimulus grids and response maps illustrate the spatial receptive-field and feature-tuning measurements obtained from the recordings. **B)** Two-photon calcium imaging sampled 12 locations distributed across V1 in each of six mice, with each location imaged at two cortical depths. Visual stimuli included localized Gabors for spatial receptive-field mapping and full-field drifting gratings for characterization of orientation, spatial-frequency, and temporal-frequency tuning.

Electrophysiological data were collected for eight mice, each with four probes targeting different regions of V1 (Figure 4A). Two-photon recordings from 12 fields of view (Figure 4B), each sampled at two depths, were recorded in six mice. Visual stimuli were designed to map receptive-field location and characterize orientation, spatial-frequency, and temporal-frequency tuning. The complete dataset and preliminary analysis code are released with this study to enable independent analyses of functional organization across V1 and tests of the unified-V2 hypothesis.

As an initial application of this resource, we used the electrophysiology dataset to ask whether hierarchy-associated functional differences among conventionally defined HVAs exceed the variation observed within V1, an established single cortical area (Figure 5). Under the unified-V2 hypothesis, these HVAs would represent different regions of a single V2 occupying a common hierarchical level above V1. If, instead, conventional HVAs correspond to distinct positions along the visual cortical hierarchy, we would expect variation among HVAs in hierarchy-associated response properties to exceed the regional heterogeneity found within V1. This analysis therefore provides a quantitative within-area benchmark for interpreting functional differences among HVAs, rather than a standalone criterion for defining areal boundaries.

**Figure 5.**
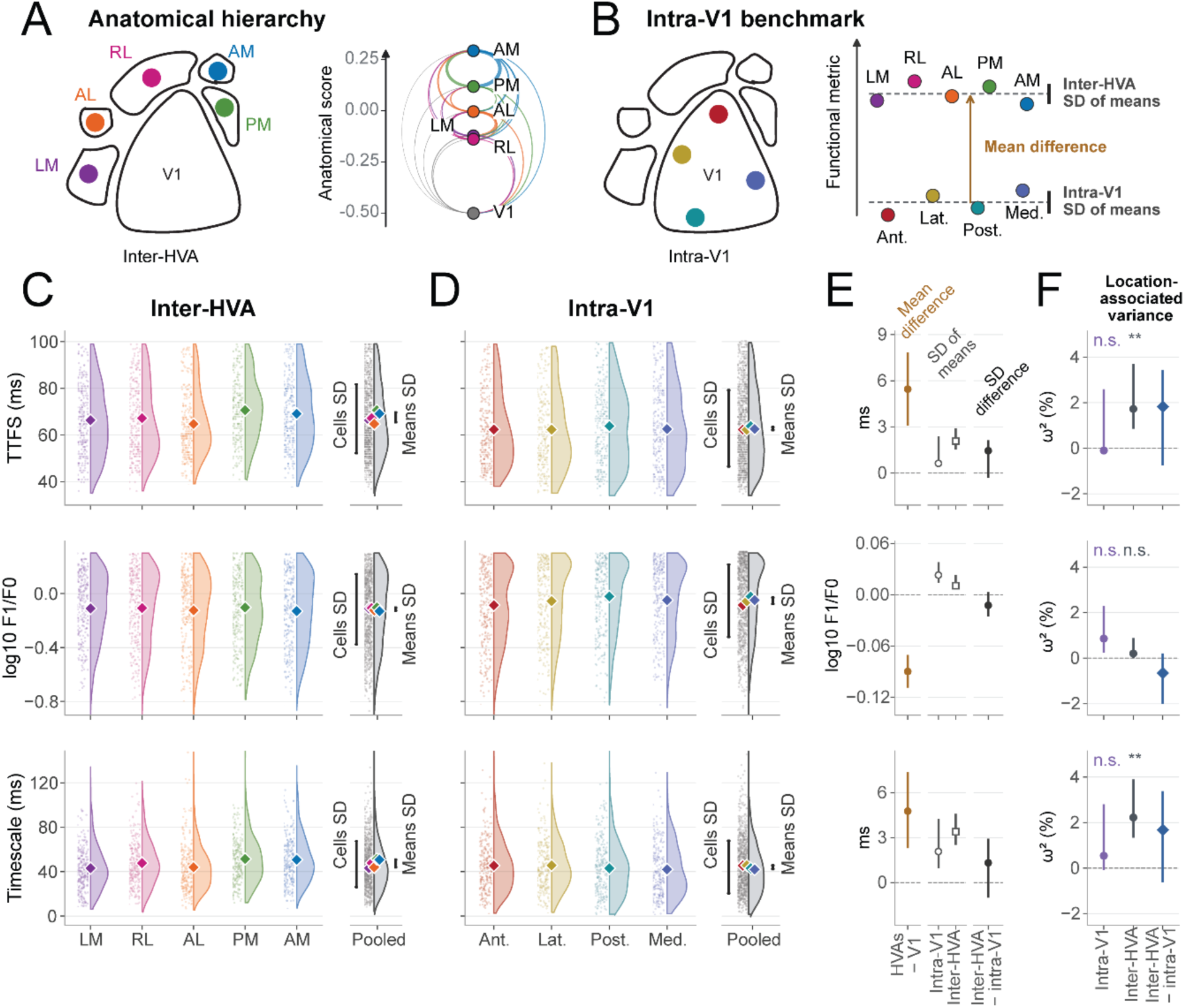
Large-scale V1 recordings provide a within-area benchmark for functional heterogeneity. **A)** Anatomical-hierarchy schematic (right) adapted from Harris et al. (2019) showing the published scores for V1 and 5 HVAs studied by Siegle et al. (2021). **B)** The four spatially separated V1 sampling regions in the new electrophysiology dataset give us a within-area benchmark for variation among HVA means (right). Cortical outlines, sampling markers and the schematic responses are illustrative. **C–D)** Single-neuron time to first spike (TTFS), phase modulation (log10 F1/F0), and neuronal response decay timescale across **(C)** the five higher visual areas (LM, RL, AL, PM, and AM) in the Allen Brain Observatory Visual Coding–Neuropixels dataset and **(D)** four V1 sampling locations in the electrophysiology dataset. Dots show subsamples of eligible neurons and half-violins show their distributions. Colored diamonds indicate session-adjusted group means, weighted to balance contributions across categories, animals and recordings. Within each population, means were recentered for display without changing their differences or SD (Methods). Pooled plots (grey, right) show distributions of individual-neuron values and fitted regional means. Bars span ±1 SD around the pooled-cell mean (“Cells SD”) and the average regional mean (“Means SD”). **E)** Point estimates with 95% confidence intervals from 5000 whole-animal bootstrapped resamples. Left: pooled-HVA-minus-V1 mean differences from simultaneous Visual Coding recordings, serving as a positive control. Middle: SDs of regional means within V1 (open circles) and across HVAs (Visual Coding; open squares). Right: inter-HVA standard deviation minus intra-V1 standard deviation. For timescale, D shows one V1 subsampling draw; V1 based estimates in E–F incorporate all ten draws (Methods). **F)** Location-associated variance estimated using bias-corrected omega-squared (ω²): intra-V1 (purple circles), inter-HVA (dark-gray circles), and inter-HVA minus intra-V1 (blue diamonds). Component estimates are percentages of total neuronal variance; differences are expressed in percentage points. Whiskers show pointwise 95% whole-animal bootstrap intervals. Component significance labels derive from one-sided within-session block-label permutation tests, Bonferroni-adjusted across six comparisons: **, adjusted p <.01; n.s., adjusted p ≥.05. Across-HVA-minus-within-V1 differences are evaluated using bootstrap intervals.

We examined three of the four response properties previously used to define a functional visual hierarchy in the mouse: response latency (time to first spike, TTFS), phase modulation (log10 F1/F0), and response decay timescale (Siegle et al., 2021). We did not examine the fourth response property, receptive field size, as our measurements did not produce reliable estimates of this feature. We quantified them using implementations adapted to improve comparability between datasets (Methods). For the within-V1 benchmark, recordings were grouped into anterior, lateral, posterior, and medial locations (Figure 5B). For the across-HVA comparison, we analyzed recordings from LM, RL, AL, PM, and AM in the Allen Brain Observatory Visual Coding–Neuropixels dataset used by Siegle et al. (2021) (Figure 5A). Hereafter, “region” refers either to one of the four sampled V1 locations or to one of the five conventionally named HVAs.

We first tested whether these measures could detect the expected broad distinction between V1 and higher visual cortex. Using Visual Coding dataset sessions containing simultaneous recordings from V1 and HVAs, the pooled HVA population had longer TTFS than V1 by 5.45 ms (95% bootstrap interval, 3.07–7.85 ms), longer response timescales by 4.77 ms (2.31–7.37 ms), and weaker phase modulation (Δlog10 F1/F0 = −0.0898; −0.1087 to −0.0705) (Figure 5E, left). Thus, all three measures distinguished V1 from higher visual cortex under matched recording conditions. This positive control confirms that the selected metrics capture the broad functional separation expected between V1 and the next level of visual cortical processing, regardless of whether higher visual cortex is treated as one area or several.

Having established this sensitivity, we next asked whether the differences among individual HVAs were larger than regional differences within V1. For each response measure, we estimated session-adjusted means for each region and quantified the spread among these means using their standard deviation (SD). We then directly compared across-HVA and within-V1 heterogeneity by subtracting the within-V1 SD from the across-HVA SD; positive values therefore indicate greater variation across HVAs. The across-HVA-minus-within-V1 differences were 1.45 ms for TTFS (95% bootstrap interval, −0.30 to 2.13 ms), −0.0124 for phase modulation (−0.0257 to 0.0031 log10 F1/F0 units), and 1.32 ms for response timescale (−1.01 to 2.93 ms) (Figure 5E, right). All three intervals included zero. Thus, at the level of regional mean responses, differences among HVAs were not clearly greater than differences among sampled locations within V1.

We then asked the same question at the level of individual neurons. Regional means alone do not indicate how strongly populations are segregated: even relatively large differences among means may account for little of the total variation when neuronal responses within each region are broadly distributed. Indeed, single-neuron distributions were broad and strongly overlapping across both V1 locations and HVAs (Figure 5C–D). We therefore quantified the fraction of neuronal variance associated with region identity using bias-corrected omega-squared (ω²; Methods). Within V1, recording location did not explain a significant fraction of variation in TTFS, phase modulation, or response timescale (Bonferroni-adjusted p = 1.000, 0.306, and 1.000, respectively). Across HVAs, HVA identity explained significant variation in TTFS and response timescale, but not phase modulation (adjusted p = 0.0024, 0.0012, and 0.678, respectively).

However, significance within one population and nonsignificance within another do not establish that the two effect sizes differ. The relevant test is therefore the direct comparison between them. The inter-HVA-minus-intra-V1 differences in ω² were 1.82 percentage points for TTFS (95% bootstrap interval, −0.74 to 3.44), −0.65 for phase modulation (−2.00 to 0.21), and 1.68 for response timescale (−0.62 to 3.39). All three intervals included zero (Figure 5F). Thus, although HVA identity accounted for detectable neuronal variation in two of the three measures, its contribution was not clearly greater than that associated with sampling different locations within V1 for all three measures.

Together, the above analyses provide a consistent picture. The selected response properties clearly distinguish V1 from higher visual cortex as broad groups, demonstrating sensitivity to the major hierarchical transition between them. Within higher visual cortex, however, neither the spread of regional means nor the fraction of single-neuron variance associated with region was significantly greater across HVAs than across locations within V1. These results do not imply that HVAs are functionally identical, nor do they establish that higher visual cortex is a single area. Rather, for the three hierarchy-associated properties examined here, the magnitude of functional differentiation among HVAs does not clearly exceed the heterogeneity observed within an established single cortical area. The results are therefore compatible with the possibility that conventionally defined HVAs represent functionally heterogeneous subdivisions of a complex V2.

This conclusion complements the retinotopic analysis above. Figure 3 showed that many reported functional differences across higher visual cortex are structured by visual field position; the present analysis shows that, for three additional hierarchy-associated properties, regional differentiation does not clearly exceed the within-area benchmark provided by V1. Neither result excludes additional, retinotopy-independent specialization among HVAs. Anatomical and connectional differences could provide such independent evidence, and we consider those next.

### Anatomical differences may reflect both areal boundaries and within-area organization

Anatomical variations provide important evidence for cortical area boundaries, but they can also reflect organization within a single area. Mouse HVAs differ in cytoarchitectonic and molecular features, projection and callosal patterns, and laminar connectivity. Each of these differences may support areal distinctions, particularly when it forms a reproducible transition that coincides with independent evidence from other modalities. However, single visual areas can also contain substantial variation in anatomical and connectional feature differences, in addition to the previously discussed variation in retinotopic and functional feature differences. Anatomical differences are therefore most informative for areal delineation when they cannot be readily explained by the underlying retinotopic map or by known forms of within-area organization. Below, we consider the major classes of anatomical evidence for mouse HVA boundaries from this perspective.

Early parcellations of mouse visual cortex used cytoarchitecture and retinotopy to distinguish two subdivisions of a single V2 (Glickfeld & Olsen, 2017). Subsequent studies identified anatomical distinctions among higher visual areas, including a prominent transition in m2AChR expression and cytoarchitecture at the LM–AL boundary (Wang et al., 2011). This transition coincides with a retinotopic reversal and provides some of the strongest convergent evidence for a conventional HVA boundary. However, molecular and cytoarchitectonic differences also occur within established areas. For example, m2AChR expression varies within V1 and is associated with functionally distinct circuits (Ji et al., 2015; Meier et al., 2025). Thus, anatomical differences provide the strongest evidence for areal boundaries when they coincide with independent transitions in other properties. Subsequent parcellations also used the spatial organization of V1 projections and callosal connections to identify reproducible subdivisions of higher visual cortex (Olavarria & Montero, 1989; Wang & Burkhalter, 2007; Wang et al., 2011). However, spatially segregated projection zones do not necessarily imply separate cortical areas. As illustrated by our simulations (Figure 2), a single area with complex retinotopy can generate multiple projection patches. Similarly, callosal patterns constrain the organization of retinotopic maps but do not necessarily distinguish internal subdivisions from areal boundaries.

Connectivity studies also reveal functional specialization across HVAs that cannot be explained solely by retinotopic sampling. Glickfeld et al. (2013) showed that V1 axons targeting different HVAs exhibit distinct visual response properties, providing evidence for selective routing of functionally specialized inputs. Other connectional differences are themselves organized by retinotopy: V1 projection targets depend partly on the location of projecting neurons (Y. Han et al., 2018), and feedback projections from HVAs to V1 exhibit systematic retinotopic biases (Morimoto et al., 2021). Importantly, neither form of specialization necessarily implies separate areas. The stripes of primate V2 receive distinct inputs from V1 (Livingstone & Hubel, 1984, 1987), while interdigitated modules within mouse V1 participate in functionally distinct cortical networks (Meier et al., 2025). Therefore, functional and connectional specialization can exist within a single visual area.

Functional specialization among V2 modules, and their relationship to retinotopy, may also arise from inputs outside V1. In particular, the superior colliculus and lateral posterior nucleus provide a retinotopically organized pathway to higher visual cortex that could contribute to differences in response properties among HVAs (Bennett et al., 2019; Calzoni et al., 2026). Consistent with a contribution from this pathway, differences in velocity preference among HVAs can persist following V1 ablation but disappear after SC lesions (Tohmi et al., 2014). Moreover, LP inputs to retinotopically matched cortical locations vary gradually around the V1 border (Juavinett et al., 2020). These findings suggest that subcortical pathways may contribute to functional differences among HVAs, including differences organized by visual field position, without necessarily defining separate cortical areas.

### Complex retinotopy beyond V1 is a common feature of secondary visual cortex in multiple species

The visual cortex immediately adjacent to V1 of several species contains complex retinotopy with multiple reversals, that resemble those used to delineate the mouse HVAs. But these reversals are often incorporated within a single area V2 rather than treated as evidence of areal boundaries. These cross-species comparisons do not establish that mouse higher visual cortex must share the same organization. Rather, they demonstrate that reversals and functional modularity need to be interpreted with coincident anatomical and cytoarchitectural evidence.

In tree shrews (Figure 6B), V2 contains several visual field reversals between alternating modules that represent central and peripheral visual field. The rationale for a single area V2 delineation includes cytoarchitecture, projections from V1, and cross-hemispheric projections (Kaas et al., 1972; Sesma et al., 1984; Wong & Kaas, 2009). In addition, despite the modular appearance of the retinotopy map, the visual field is represented in V2 by a single continuous transformation, with azimuth following a complex progression and elevation a smooth one, such that the reversals are folds within one map rather than separate maps (Sedigh-Sarvestani et al., 2021). Functional property differences observed in V2 stripes are also related to their different retinotopic biases (Sedigh-Sarvestani et al., 2021). Ferret visual cortex similarly contains complex retinotopic maps with internal reversals (Figure 6C), but these reversals are distinguished from boundaries between areas 18 and 19 using convergent anatomical and functional evidence (Dell et al., 2019; Innocenti et al., 2002; Manger et al., 2002). As in tree shrew V2, the extrastriate cortex in ferrets exhibits complex internal retinotopy, with individual modules representing restricted portions of visual space (Manger et al., 2002). Related complexity has also been reported in cat visual cortex (Mallot, 1985; Tusa et al., 1979). Modeling work proposed that such complex retinotopy can arise during development as a consequence of the shape of the extrastriate cortical sheet, and that multiple stable solutions could account for inter-individual variability in the cat, as well as other species (Wolf et al., 1994).

**Figure 6.**
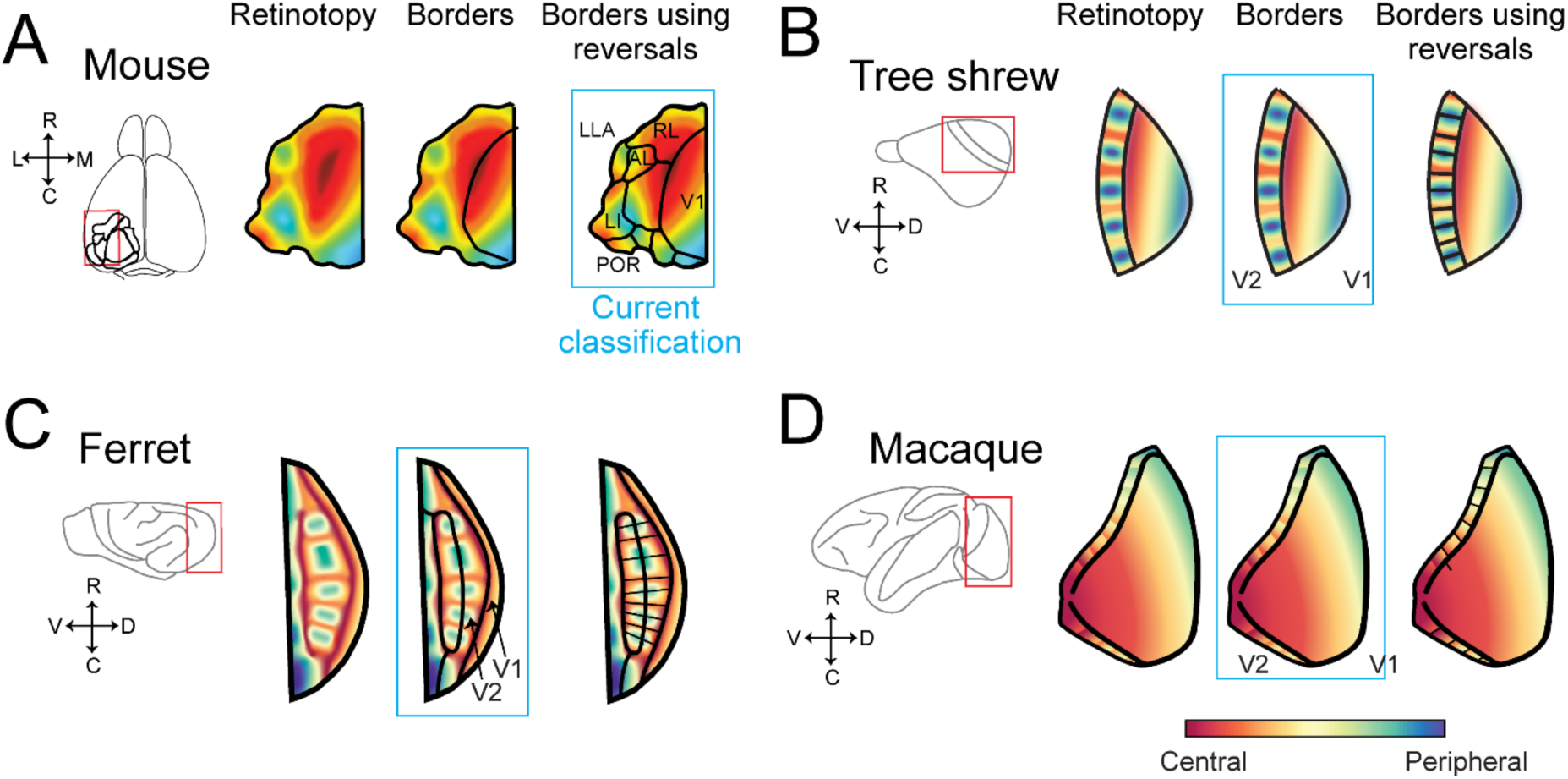
Complex retinotopy beyond V1 is a common feature of secondary visual cortex in multiple species. **A)** Retinotopy maps in mouse visual cortex, schematics created based on data from (Zhuang et al., 2017). Hypothetical borders using reversals alone are shown on the right. These borders are similar to the currently accepted area delineations shown in the blue box. **B)** Retinotopic maps in tree shrew visual cortex. Data from (Sedigh-Sarvestani et al., 2021). **C)** Retinotopic maps in ferret visual cortex. Data from (Manger et al., 2002). **D)** Schematic of retinotopic maps in the visible portion of macaque V1 and V2. Data from (Roe & Ts’o, 1995; Shipp & Zeki, 2002b). Color maps are different for each species and are simply meant to indicate relatively more central or peripheral regions of the visual field.

A series of elegant papers in the 1980s and 1990s showed a modular, striped, organization in macaque V2 (Hubel & Livingstone, 1987; Roe & Ts’o, 1995; Shipp & Zeki, 2002a). The stripes exhibited distinct functional feature preferences, including for color, motion, and horizontal disparity. The underlying retinotopic map has been less clearly defined, in part due to the lack of optical imaging measurements in macaque V2. Published reports based on electrode penetrations across V2 indicate multiple reversals between relatively more foveal or peripheral regions of the visual field (Roe & Ts’o, 1995; Shipp & Zeki, 2002b). Regardless of the precise geometry of these internal retinotopic transformations, macaque V2 provides a clear example in which pronounced functional modularity—encompassing differences in color, motion, and disparity processing— exists within a single established cortical area. These functional subdivisions are therefore treated as modules of V2 rather than independent cortical areas.

Comparisons within rodents are particularly informative because the organization proposed for mouse visual cortex is not universal across the order. In the Eastern gray squirrel, electrophysiological mapping identified a single V2 (area 18) bordering V1, with a systematic representation of the contralateral visual hemifield that approximately mirrors V1 (Hall et al., 1971). Subsequent anatomical and connectional studies supported this organization while also revealing substantial modular structure within V2 (Gould, 1984; Kaas et al., 1989). Outside the squirrel lineage, the visually specialized agouti likewise has an architectonically and physiologically defined V2 bordering V1, with an orderly representation of the contralateral visual field (Elston et al., 2006; Picanço-Diniz et al., 1989). The squirrel-related lineage is especially informative phylogenetically: molecular analyses place it near the base of Rodentia, as sister to the mouse-related and Ctenohystrican clades (Churakov et al., 2010). Thus, the presence of a single V2 in squirrels is consistent with a V2-centered organization being ancestral within rodents, whereas the mouse is a more derived lineage. Both squirrels and agoutis are also more visually specialized than mice. It would therefore be surprising if the nocturnal mouse, with its smaller visual cortex, had independently evolved a more elaborate parcellation of extrastriate cortex than these diurnal, highly visual relatives (Laramée & Boire, 2014; Rosa & Krubitzer, 1999). A more parsimonious possibility is that mouse extrastriate cortex retains the broader V2-centered organization seen in other rodents, with some conventionally defined HVAs representing modules within it.

The comparisons above do not imply that secondary visual cortex must follow an identical blueprint across mammals; species-specific factors such as evolutionary lineage, visual field geometry, cortical sheet geometry and binocular overlap may influence map organization. The complex V2 map of the lateral-eyed tree shrews however, suggests that strong binocularity is not a prerequisite for internal retinotopic reversals. In summary, complex and modular retinotopy beyond V1 occurs in several evolutionarily distant mammals, but the resulting modules are not uniformly treated as independent areas. These comparisons demonstrate that field-sign reversals, redundant representations, and functional modularity must be interpreted in the context of other anatomical, connectional, and functional evidence. They therefore motivate reconsideration of the criteria applied to mouse higher visual cortex.

## DISCUSSION

Higher visual areas in the mouse are delineated by retinotopic reversals, by anatomical and functional differences among the resulting regions, and by a hierarchical ordering built on such differences. Here we show that several features used to distinguish mouse HVAs can also arise within a single cortical area, motivating reconsideration of the conventional parcellation. Our simulations demonstrate that a single complex retinotopic map can generate multiple field-sign reversals, redundant representations, and spatially segregated projection zones (Figure 2). Many functional differences among conventionally defined HVAs covary with their systematic biases in visual field representation (Figure 3). Finally, variation in the response properties used to order HVAs hierarchically did not significantly exceed variation across sampled locations within V1 in our newly collected within-V1 dataset (Figures 4–5). Together with cross-species evidence for modular organization within established visual areas (Figure 6), these findings motivate the unified-V2 model in which many conventional mouse HVAs represent functionally distinct subdivisions of a larger V2.

This possibility has a long history. Malach (1989) and Kaas et al. (1989) proposed that multiple extrastriate projection zones in rodents could represent modules of a single V2. The theoretical work of Wolf et al. (1994) offered a possible mechanism for such complex retinotopic organization: a single elongated visual area could develop a complex retinotopic map with multiple modules rather than one smooth map. Rosa and Krubitzer (1999) later framed the question as a choice between a “simple” hypothesis, in which V1 is bordered laterally by a single V2, and a “complex” hypothesis, in which V1 is bordered by multiple areas. They argued that comparative evidence clearly favors the simple hypothesis. Krubitzer and Huffman (2000) suggested that cortical modules may represent intermediate stages in the emergence of new cortical areas, an idea later reviewed in the context of mouse and squirrel visual cortex (Laramée & Boire, 2014). Striedter and Northcutt (2020) noted that the unusual spatial arrangement of the numerous proposed mouse visual areas, relative to primates, leaves open the possibility that some are modules rather than independent areas. Kaas et al. (2021) likewise highlighted an alternative interpretation in which the cortex bordering V1 in rats and mice comprises just two areas, V2 and a medially located prostriata. Our proposal therefore revisits a longstanding alternative. Our contribution is to revisit this alternative using modern retinotopic evidence, simulations constrained by mouse cortical geometry, and large-scale functional recordings.

The central question here is what evidence is sufficient to distinguish an areal boundary from organization within an area (Petersen et al., 2024; Rosa, 2002). We argue that boundaries are most strongly supported when independent features, including topography, architectonics, connectivity, and function, show reproducible and spatially coincident transitions. Such evidence is not uniform across conventional HVA boundaries. The LM–AL border, for example, combines a clear molecular and cytoarchitectonic transition with a retinotopic reversal (Wang et al., 2011), providing strong evidence for an area boundary. Other regions, including RL, exhibit distinctive functional properties (La Chioma et al., 2019; Y. Zhang et al., 2026), but these functional specializations do not necessarily coincide with comparable anatomical transitions. This lack of convergence is compatible with a larger area containing specialized modules whose strongest distinctions need not define common areal borders. However, the available data also leaves room for an intermediate organization in which some HVAs are subdivisions of V2 while others remain independent areas.

Anatomical and functional hierarchy provide another means of evaluating these alternatives. Laminar projection patterns distinguish V1 from higher visual cortex, whereas connections among HVAs frequently resemble lateral pathways (D’Souza et al., 2022). Although connectivity-based analyses assign distinct hierarchical positions to HVAs (Harris et al., 2019), functional measurements do not consistently follow these orderings. For example, Akella et al. (2025) found that RL departed from hierarchy-associated trends in neuronal variability and stimulus encoding, with significant correlations emerging only after excluding RL. Our analysis based on the new largescale V1 dataset provides a complementary within-area benchmark: across three hierarchy-associated response properties, variation among HVAs did not significantly exceed variation across sampled locations within V1. These results do not establish equivalence between HVAs, particularly given differences in sampling between datasets, but suggest that current evidence distinguishes V1 from higher visual cortex more clearly than it establishes separate hierarchical stages among individual HVAs.

The unified-V2 and multi-area models make distinguishable predictions. Distinct HVAs would be supported by reproducible borders at which multiple independent properties change together—retinotopy-controlled functional discontinuities, coincident structural transitions, abrupt connectivity changes at matched receptive-field locations, or hierarchical discontinuities—and by independently organized visual field maps within individual HVAs. Conversely, unified V2 would be favored by cellular-resolution evidence for a continuous complex retinotopic transformation across conventional borders, functional and connectional properties better predicted by retinotopic position than HVA identity, and inter-HVA differences comparable to within-area heterogeneity. The V1 dataset and analysis tools are released to enable independent tests of these predictions, including analyses that challenge our interpretation.

The unified-V2 model also changes how we might think about cortex beyond this region. If the cortex immediately surrounding V1 is instead treated as V2, this also raises the question of where later stages of mouse visual processing lie. POR is a particularly interesting target because of its position in posterior temporal cortex, strong SC–LP-dependent visual drive, and sensitivity to moving objects (Beltramo & Scanziani, 2019; Brenner et al., 2023). This combination is notable in light of the proposal that primate MT emerged from an ancestral temporal cortical territory driven by the SC–pulvinar pathway (Kaas et al., 2021).

Ultimately, area definitions are necessarily operational as useful conventions for describing biological organization. More than the particular nomenclature ultimately adopted, it is important that our conventions are continually tested against new data. We present the unified-V2 model as one testable alternative to the conventional mouse HVA parcellation, not as an established reclassification. By making our predictions explicit and releasing the V1 benchmark dataset, we hope to encourage independent tests of these models and a more consistent application of areal delineation criteria across species.

## MATERIALS AND METHODS

### Model geometry and visual field extent (**Figure 2**)

To apply the modeling techniques used in previous papers, we first needed to account for key differences in anatomy of the mouse visual cortex, including the ring-like shape of the cortical sheet surrounding V1 and the extent and geometry of the mouse visual field represented in V1 (Figure 2A). Because the visual field extent sampled by widefield imaging is restricted, particularly in elevation, we modeled a broader visual field envelope spanning approximately 0–140° azimuth and −40–60° elevation, constrained by electrophysiological measurements from Wagor et al. (1980) and the rounded visual field coverage reported by Garrett et al. (2014). The initial cortical footprint was derived from the population retinotopic maps of Waters et al. (2019). For each animal, we measured the contiguous extrastriate extent along outward normals from the V1 border. Measurements from repeated sessions were first averaged within mouse, and the median extent across mice was projected along corresponding normals from a common V1 contour. This initial footprint was expanded to include selected extrastriate support regions, each thresholded at 50% of its own peak across-mouse support. These included an inferred LI-like region; RLL was excluded from the expansion masks. The original footprint was retained, and the combined envelope underwent morphological closing with a 50-µm radius and hole filling.

We modified the algorithm to weight the interactions between neighbors by the inverse square of their distance in cortical space to account for unevenly spaced nodes. In addition, we also expected a lower cost associated with the smoothness constraint due to larger receptive fields in the mouse visual system. The mouse’s smaller eyes result in low spatial resolution as the visual field is projected through the pupil onto a smaller retinal surface. In V1 this results in much larger receptive fields and beyond this, allows for bigger steps in the visual field to be taken by adjacent regions of cortex while preserving an overlapping representation. Given this, we expected that a lower weight on the smoothness constraint in our model may result in a better representation of the higher order visual areas’ retinotopy.

### Model fitting

The surrounding cortical domain comprised a grid of 160 × 16 trainable nodes, periodic around V1, with 160 additional fixed border samples. We prescribed the correspondence between cortical border positions and visual-field locations independently of the fitted V1 reference map. Both fits used this boundary correspondence and the same set of 2,000 visual-field targets. Targets were sampled using a golden-angle spiral with density parameter c = −0.05, producing approximately uniform areal sampling with weak weighting toward the sampling center at 0° azimuth and 20° elevation. The objective combined visual-field coverage, local smoothness weighted by inverse squared cortical distance, a penalty for deviations from the prescribed border RFs, and local metric isotropy:

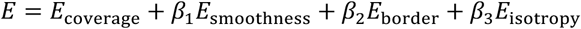

The coefficients were *β*_1_ = 1.45 *x* 10^-5^, *β*_2_ = 100, and *β*_3_= 0.030. Optimization used RMSProp over 3,000 iterations, an initial learning rate of 0.1, momentum of 0.8, and an initial annealing width of 44°. The V1 reference used a nonperiodic 41 × 41 grid, comprising 1,521 trainable interior nodes and 160 fixed perimeter nodes, with matched objective coefficients and optimization schedules.

### Visualization and subdivision

In Figure 2B, V1 RF coordinates were interpolated before computing field sign from the normalized azimuth–elevation Jacobian, followed by 0.075-mm field-sign smoothing. The surrounding-domain field-sign map used the native-mesh calculation and 0.10-mm smoothing employed for subdivision. The meshes in Figure 2A and C received additional display interpolation and light coordinate smoothing, with boundary restoration. In the middle view of Figure 2C, surrounding RF locations were mapped to corresponding cortical positions through the fitted V1 triangular mesh, with piecewise cubic Hermite interpolation used to complete missing or ambiguous inverse correspondences. Height was an illustrative coordinate based on cortical distance from the V1 border. Retinotopic subdivisions were obtained using an adaptation of Garrett et al. (2014), with 0.10-mm field-sign smoothing, a sign threshold of 0.25, 0.05-mm eccentricity smoothing, a merge-overlap threshold of 0.05, and a redundancy threshold of 1.01. Before partitioning, fitted surrounding RF coordinates were recentered and independently rescaled to span 80 units in azimuth and 50 units in elevation. This rescaling was introduced to accommodate the model’s broader visual-field extent when applying the adapted Garrett subdivision procedure. Eccentricity and visual-coverage calculations used these rescaled coordinates; visual-field plots retain the original degree coordinates. No internal subdivision boundaries or subdivision count were prescribed during model fitting.

### Simulated tracer mapping

Three simulated injection disks, each of radius 0.19 mm, were placed within the fitted V1 map. Correspondences were calculated using piecewise-linear interpolation over the V1 triangular mesh. A surrounding node was labeled if any corresponding V1 position lay within an injection disk. Colored meshes connect labeled nodes that are adjacent in cortex. Predicted labeling therefore reflects retinotopic correspondence.

### Retinotopic registration and spatial-and temporal-frequency analysis (**Figure 3F–G**)

We reanalyzed the Han et al. (2022) dataset using the reliable-cell flags and fitted tuning parameters supplied with the data. Two-photon cell coordinates were aligned to each animal’s widefield map and then to a common cortical map. Digitized azimuth and elevation maps from Zhuang et al. (2017) were aligned to this common map using an affine transformation, and each cell was assigned the azimuth and elevation at its registered cortical location. These values therefore represent registration-based estimates of retinotopic position rather than independently measured single-cell receptive-field centers. Azimuth was limited to 0–100° and elevation to −40–40°. Eccentricity from the focea was calculated as 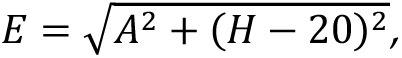 where *A*is azimuth and *H*is elevation in degrees, corresponding to a reference point at 0° azimuth and +20° elevation. Analyses included cells assigned to LM, AL, RL, AM, PM, or LI with finite tuning and retinotopic coordinates. The combined POR/P population was excluded because these areas were not separated in the source dataset. A total of 15,254 cells contributed to each plane fit.

Spatial and temporal frequencies were expressed in octaves as *z_SF_* = log_2_(*SF*/0.02)and *z_TF_* = log_2_(*TF*/0.5), respectively. For each metric, we fit a plane *μ* = *aE* + *bH* + *c*across individual cells. Values at or below 0.01 octaves and at or above 3.99 octaves were treated as lower-and upper-censored, respectively, relative to nominal limits of 0 and 4 octaves. Censored values were incorporated using an iterative Gaussian-tail imputation and least-squares fitting procedure in which the plane and residual scale were updated until the scale changed by less than 10^-6^or 100 iterations were reached. All cells were equally weighted. The nominal limits correspond to 0.02–0.32 cycles/degree for SF and 0.5–8 Hz for TF; Han et al. (2022) tested TF stimuli through 16 Hz.

For the displayed tuning distributions, residuals, and mouse summaries, we included cells strictly within the corresponding analysis bounds, yielding 12,105 SF observations and 10,291 TF observations. Residuals were calculated as observed minus fitted log2 frequency (*z* − *μ*), and fitted values were transformed back to physical units for display. Violin plots show kernel-density estimates of pooled cell values. Within each area, values were first averaged across cells from the same mouse; gray points show individual-mouse means, and the large white point and bar show the equally weighted mean across mice and its t-based 95% confidence interval. The numbers of contributing mice for LM, AL, RL, AM, PM, and LI were 10, 8, 8, 6, 6, and 9, respectively. Mean retinotopic-position markers were calculated from the pooled cell coordinates.

### Largescale V1 data collection (Figure 4). Electrophysiology

Mice received a headpost and a SHIELD cranial implant permitting repeated probe access (Bennett et al., 2024). Following recovery, animals underwent two weeks of handling and head-fixation habituation following established procedures (de Vries et al., 2020). Mice were gently handled for 1–2 min on days 1–2, followed by handling and 5–10 min of head fixation on days 3–5. On days 6–10, head-fixed visual stimulation increased progressively **from** 10 minutes to 50 minutes across 5 days. Animals were additionally habituated to the Neuropixels rig for one week, including three sessions in increasing time increments with the last habituation being comparable in length to the recording sessions. Visual cortical areas were delineated using intrinsic signal imaging (ISI) under light isoflurane anesthesia (1–1.4%). Contrast-reversing checkerboard bars swept across the visual field in four cardinal directions at 0.1 Hz. Fourier-derived azimuth and elevation maps were combined into visual field sign maps, averaged across at least three acquisitions, and registered to cortical vascular landmarks (Garrett et al., 2014).

Recordings were performed in awake, head-fixed mice allowed to locomote on a freely rotating disc, using established multiprobe recording procedures (Durand et al., 2023). Up to six Neuropixels 1.0 probes were inserted to 3.1 mm at 200 µm/min, retracted by 100 µm to reduce tissue compression, and allowed to settle for approximately 30 min. The 384 recording sites nearest each probe tip were sampled at 30 kHz for action potentials and 2.5 kHz for local field potentials using Open Ephys. Visual stimuli were presented on a gamma-corrected, 60-Hz monitor positioned 15 cm from the right eye. Eye, face, and body videos were acquired at 60 Hz, and locomotion was monitored with a rotary encoder. Photodiode signals and digital synchronization pulses aligned stimulus presentation and behavioral measurements with neural activity. Spikes were sorted using Kilosort 4 (Pachitariu et al., 2024) and subjected to automated quality assessment. Recording locations were assigned using fluorescent probe-track reconstruction and registration to the Allen Common Coordinate Framework (Wang et al., 2020). For this, we used the IBL ephys alignment GUI (https://github.com/AllenNeuralDynamics/ibl-ephys-alignment-gui). Based on firing rate, LFP power, as well as spike cross-correlograms, we placed reference lines delineating borders of areas. Through this procedure, we consistently aligned the top of the cortex (Layers 2/3, where we expect spiking activity) using cross-correlograms. Other alignment locations such as white matter tracts, thalamus, and subcortical regions are only made when there are clear electrophysiological landmarks (i.e. a stark increase in firing rate for thalamus, a stark decrease in activity in white matter, etc). If features such as these are not present, the GUI’s probe track interpolation is accepted. Notably, this interpolation is used for cortical layer assignments between Layers 2/3 and the bottom of cortex.

### Two-Photon Imaging

Mice expressing GCaMP6f received a headpost and a chronic 5-mm cranial window over visual cortex using established surgical procedures (Groblewski et al., 2020). Following recovery, animals completed the handling and progressively extended head-fixation habituation described above, followed by a 30-min acclimation session on the mesoscope without visual stimulation. ISI-derived retinotopic maps were registered to live epifluorescence images using superficial vascular landmarks to identify the targeted visual areas. The objective was aligned perpendicular to the cranial window, and stage coordinates, surface images, and cellular reference images were retained to support repeated targeting of the same fields and neurons across sessions.

ISI-derived retinotopic maps were used to identify functionally defined visual areas and target locations for two-photon imaging. On the first imaging day, the ISI targeting map was registered to a high-resolution reference image of the cortical surface vasculature acquired through the cranial window. Superficial vascular landmarks, including vessel branching patterns and intersections, were used to align the ISI map with live epifluorescence images and thereby transfer the retinotopically defined target location to the cranial window. The aligned target was then localized under two-photon imaging by comparing the live 2P surface image with the registered vascular reference and ISI map. A 2P reference image of the cortical surface was acquired for each imaging site, and stage coordinates and reference images were retained to facilitate consistent targeting of the same cortical locations across subsequent imaging sessions.

Calcium imaging was performed in awake, head-fixed mice on a freely rotating disc using a dual-beam mesoscope based on the published 2P-RAM architecture (Sofroniew et al., 2016) and controlled by customized ScanImage software. Excitation was provided at 910 nm, and individual imaging fields measured 400 × 400 µm, sampled at 512 × 512 pixels and approximately 11 Hz per plane. Imaging planes sampled cortical depths spanning superficial cortex through layer V, with laser power adjusted according to depth and fluorescence intensity. Reference z-stacks extending 30 µm above and below each imaging plane, in 0.75-µm steps, supported assessment of axial stability. Imaging was synchronized with visual stimulation, locomotion, and eye and behavioral videos acquired at 30 Hz. Processing included nonrigid motion correction, correction of inter-plane fluorescence crosstalk, cellular segmentation, neuropil subtraction, and calculation of ΔF/F traces. Deconvolved activity was estimated using OASIS (Friedrich et al., 2017). Sessions were assessed for saturation, photobleaching, motion, and synchronization failures.

### Example neurons from the V1 datasets (Figure 4)

For the electrophysiology (Ephys) dataset example, the signal was computed as the difference in firing rate between the baseline and evoked activity, with the baseline corresponding to the grey screen epochs of the stimulus block. For the calcium imaging dataset (Ophys), the signal was the dF/F traces directly available from the repository. To display the response properties of the example neurons, the signals were aligned to the stimulus onset and averaged over the samples in the presentation window (0.25s for Ephys and 0.32s for Ophys for Gabor patches; and 1s and 1.05s for gratings, respectively), giving one response per trial. Trials sharing a stimulus condition were averaged. Receptive fields were obtained by grouping Gabor patch trials by screen position; orientation tuning by grouping grating trials by orientation (pooled across spatial and temporal frequencies), and the spatial × temporal frequency matrix was taken at the neuron’s preferred orientation (orientation evoking the maximum average response).

### Comparison of functional variation within V1 and across HVAs (**Figure 4-5**)

The OpenScope electrophysiology dataset sampled four targeted locations within V1, designated anterior (probe A), lateral (E), posterior (C), and medial (B) in eight mice. The imaging dataset sampled 12 V1 fields of view at two imaging depths in six mice; Figure 4B shows the 12 fields from a single mouse overlaid on its widefield retinotopic map.

For the functional hierarchy comparison (Figure 5), we analyzed three electrophysiological response properties previously associated with visual cortical hierarchy (Siegle et al., 2021): time to first spike (TTFS), phase modulation (log10 F1/F0), and response decay timescale. Probe locations were registered to the Allen CCFv3 atlas and only neurons anatomically assigned to V1 were included in the functional analysis, and session-by-location groups containing fewer than five eligible neurons were excluded. For comparison across higher visual areas, we also analyzed recordings from LM, RL, AL, PM, and AM in the Allen Brain Observatory Visual Coding–Neuropixels dataset described by Siegle et al. (2021). Region was treated as a categorical variable, with four sampling regions within V1 and five conventionally named HVAs. Methods were adapted from Siegle et al. (2021) to improve comparability between datasets: we used F1/F0 in place of the spectral modulation index, restricted the drifting-grating analysis to shared stimulus conditions, and subsampled flash trials from 150 to 75 trials per polarity for the response-timescale analysis. Differences in recording conditions and neuronal sampling between datasets may nevertheless contribute to the estimated heterogeneity. The metric definitions and inclusion criteria are detailed below.

The main heterogeneity comparison used OpenScope V1 recordings to estimate variation across locations within V1 and Visual Coding recordings to estimate variation across HVAs. V1 recordings from Visual Coding were used only in a separate positive control, which compared V1 and HVA responses recorded simultaneously under matched session conditions (Figure 5E, left).

TTFS was calculated using procedures adapted from Siegle et al. (2021) on responses to full screen flashed stimuli at each neuron’s preferred flash polarity, determined by mean spike count over 0-250ms after the recorded stimulus onset. TTFS was the median first-spike latency across the preferred-polarity trials, as measured in 1ms bins from 30 through 200ms after onset; trials without spikes in this window were omitted. Inclusion was gated on a significant (one-sided paired Wilcoxon test, Benjamini–Hochberg q <.01), positive response to the preferred flash polarity with mean firing rate during 30–180ms after flash onset exceeding the mean baseline rate during −100–0ms by ≥2 Hz and a TTFS < 100ms.

Phase modulation was quantified using F1/F0 (Skottun et al., 1991), rather than the spectral modulation index, to reduce sensitivity to differences in response phase across trials. Analyses used each neuron’s preferred stimulus orientation and temporal frequency chosen from the restricted shared drifting-grating conditions: four orientations (0°, 45°, 90°, and 135°), five temporal frequencies (1,2,4,8,15 Hz) with spatial frequency fixed at 0.04 cycles per degree. F1/F0 was calculated as the amplitude of the first Fourier harmonic divided by the mean response over the first second of each trial, averaged across trials with nonzero mean responses, and log10-transformed.

Response decay timescale was calculated following Siegle et al. (2021) by fitting an exponential decay to the spike-count correlation curve over the interval 40-290ms after flash onset in 10ms bins. Inclusion required a timescale less than 300ms with a standard error less than 20ms and more than 50 spikes in the analysis window. V1 (OpenScope) sessions were subsampled down to 75 of 150 trials per polarity, across 10 draws, to match the Visual Coding trial counts.

For each measure and population, category means (probe location in V1, or HVA area identity) were estimated using weighted linear models with additive session effects to account for session-to-session shared shifts across categories. For display in Figure 5C–D, fitted regional means were shifted by a common offset within each population to align their average with the equally weighted average of the raw regional means, preserving differences among means and their SD. Categories received equal total weight, divided equally among animals within category, recordings within animal/category, and neurons within recording/category. Spread across categories was quantified as the standard deviation of the fitted means, using denominator K, where K was four for V1 locations and five for HVAs. For timescale, the 10 draw-specific category variances were averaged before taking the square root.

To quantify the cell-level variance associated with the category identity, we used a bias-corrected one-way omega-squared (Lakens, 2013), with equal neuron weights and no session adjustments,

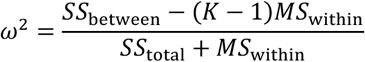

where SS_between_ and SS_total_ denote the between-group and total sums of squares, respectively; MS_within_ denotes the within-group mean square, and K is the number of groups. Negative estimates were retained, and timescale ω² estimates were averaged across the ten draws. To test whether category explained more cell-level variation within each population than expected under shuffled labels (Figure 5F), we compared the observed omega-squared estimates to those drawn from 5000 within-session block-label permutations. Category labels were reassigned within each session, keeping all neurons from the same category together, to test the null that category labels are interchangeable within V1 and across the HVAs. The one-sided permutation p-values used a plus-one correction and were Bonferroni-adjusted across the six comparisons.

To directly compare heterogeneity across the HVAs (inter-HVAs) with that within V1 (intra-V1), we subtracted the intra-V1 estimate from the inter-HVA estimate for both SD (Figure 5E) and ω² (Figure 5F), with ω² differences expressed in percentage points. Pointwise 95% percentile confidence intervals were obtained from 5,000 whole-animal bootstrap resamples. Animals were sampled independently with replacement within each dataset, retaining the original number of animals and preserving dependence among neurons and probes within animals while estimating uncertainty associated with animal sampling. In each replicate, models were refitted and both component estimates and their differences were recalculated. For response decay timescale, one trial subsample was selected for each V1 (OpenScope) session occurrence so that the draw was shared across its neurons and recording locations.

For the matched V1-HVA comparison (Figure 5E, left), the mean difference was estimated using a weighted linear model with group (V1 or pooled HVAs) membership and additive session effects. The V1 and pooled-HVA groups received equal total weight. Within each group, weight was divided equally among its constituent areas (one for V1, five for HVAs), then among animals within each area and recordings within each animal. Neurons within each area–recording combination received equal weights, but their weights could differ across recordings and areas. This balanced contributions despite unequal sampling. The analysis included 40, 28 and 50 eligible sessions for TTFS, phase modulation and response decay timescale, respectively. Sessions were not required to contain all five HVAs. Confidence intervals were estimated by resampling animals, retaining the simultaneously recorded V1 and HVA data together within each sampled animal.

## ACKNOWLEDGEMENTS

We would like to thank Isaac Rhim for sharing his insights on the retinal cone opsin gradients in mice. We thank Xu Han in the lab of Vincent Bonin for making their data and code available. We thank Marina Garrett, Mai Morimoto, and Matteo Carandini for helpful comments on an earlier version of the manuscript. Lastly, we thank Fred Wolf for insightful feedback and discussion.

The experimental dataset in this project was obtained as part of the OpenScope program, which is operated by the Allen Institute, Neural Dynamics program. We thank the OpenScope steering committee and Karel Svoboda for their support. This research was also supported by the Allen Institute, founded by Jody Allen – chair and cofounder of Allen Family Philanthropies, and the late Paul G. Allen – investor, philanthropist, and co-founder of Microsoft.

## AUTHOR CONTRIBUTIONS

Contributions of each author are listed below. This author contribution table was created with a modified version of CTAB: https://try-ctab.github.io/#/.

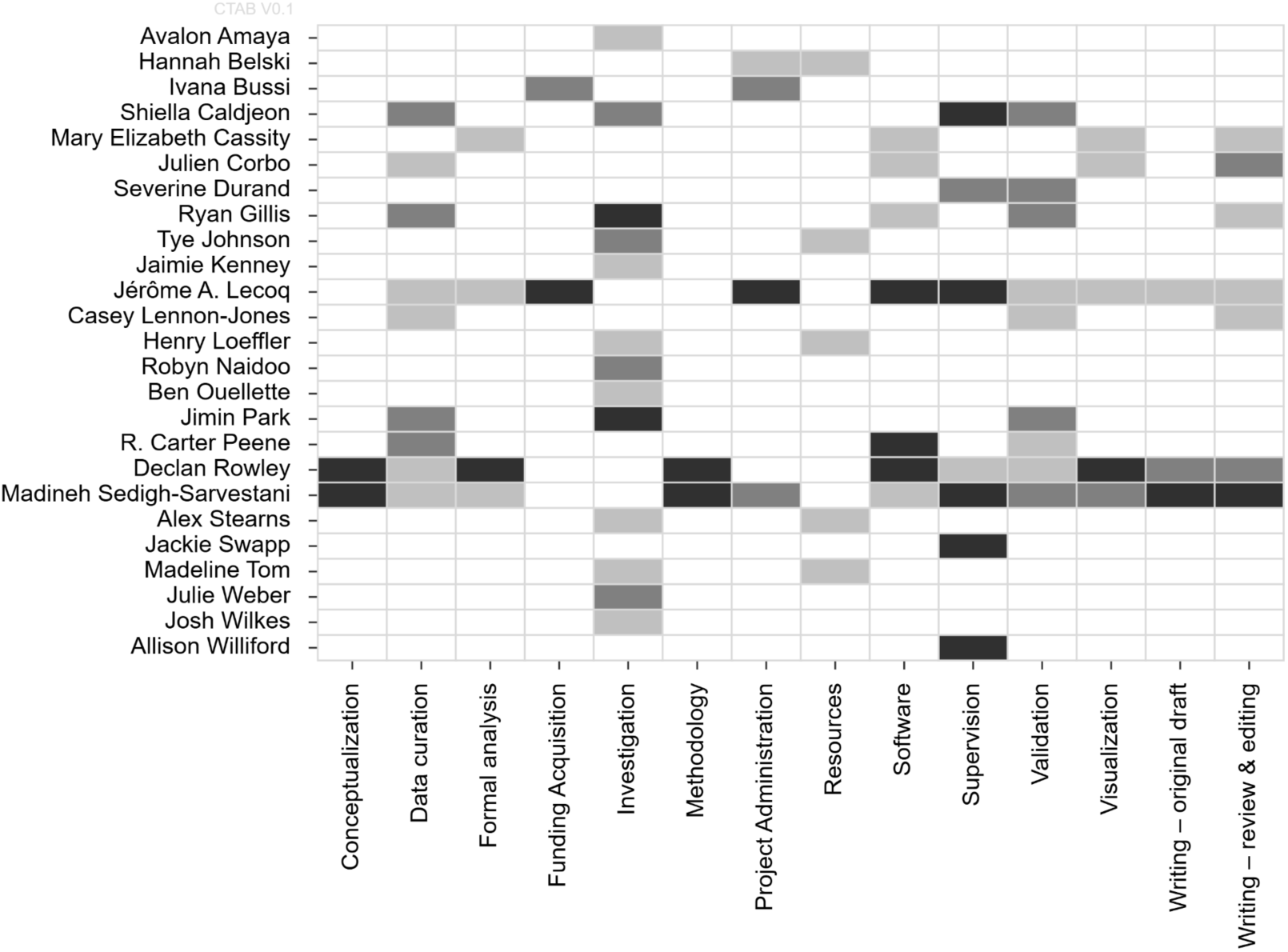

## DATA AND CODE AVAILABILITY

The Allen Brain Observatory Visual Coding–Neuropixels dataset used for the higher visual area analyses is publicly available through the Allen Institute and was originally described by Siegle et al. (2021). The new OpenScope V1 datasets reported here are deposited in DANDI Archive and made publicly available. Electrophysiology data are publicly available on the DANDI Archive (RRID:SCR_017571) at the following URL: https://dandiarchive.org/dandiset/001568. Two-photon imaging data are publicly available on the DANDI Archive (RRID:SCR_017571) at the following URL: https://dandiarchive.org/dandiset/001709. Analysis code for Figures in the paper as well as helper code to get started with data analysis of the V1 data is available at https://github.com/SarvestaniLab/MouseV2.

## AI DISCLOSURE

Generative AI tools (Claude, Claude Code, ChatGPT, GitHub Copilot) were used to assist with writing and debugging analysis and visualization code, converting code from MATLAB to Python, and to improve the text of the manuscript. All AI-assisted recommendations were reviewed, tested, and validated by the authors. All analyses, results, and interpretations were performed and verified by the authors. The authors take full responsibility for the content of the manuscript.

